# Liquid phase electron microscopy of bacterial ultrastructure

**DOI:** 10.1101/2024.02.27.580996

**Authors:** Brian J. Caffrey, Adrián Pedrazo-Tardajos, Emanuela Liberti, Ben Gaunt, Judy S. Kim, Angus I. Kirkland

**Author notes:** These authors contributed equally to this work (Brian J. Caffrey, Adrián Pedrazo-Tardajos).

## Abstract

Recent advances in liquid phase scanning transmission electron microscopy (LP-STEM) have enabled the study of dynamic biological processes at nanometre resolutions, paving the way for live-cell imaging using electron microscopy. However, this technique is often hampered by the inherent thickness of whole cell samples and damage from electron beam irradiation. These restrictions degrade image quality and resolution, impeding biological interpretation. Here we detail the use of graphene encapsulation, STEM, and energy-dispersive X-ray (EDX) spectroscopy methods to mitigate these issues, providing unprecedented levels of intracellular detail in aqueous specimens. This work demonstrates the potential of LP-STEM to examine and identify internal cellular structures in thick biological samples, in a radiation resistant, gram-positive bacterium, Deinococcus radiodurans using a variety of imaging techniques.

## Introduction

Liquid phase electron microscopy (LP-EM) is a technique that enables the analysis of intracellular, real-time dynamics with nanometre precision in native, hydrated, aqueous environments. This rapidly emerging field of research has provided unique insights into hard materials ^1^, such as the formation of platinum nanocrystals ^2^ and the electrochemical analysis of lithium-ion batteries ^3^. More recently, LP-EM has also expanded to the analysis of biological materials such as proteins ^4^, bacteria ^5^, and even eukaryotic yeast ^6^ and human cells ^7,8^. However, ultrastructural analysis of the internal components of these biological materials is difficult due to limitations in sample thickness and electron beam fluence.

Central to addressing these limitations are considerations of the most appropriate imaging geometry for LP-EM imaging of biological samples. Conventional transmission electron microscopy (TEM) is the most widely used imaging technique in biological LP-EM ^9^ and in structural biology more generally ^10^. The popularity of brightfield TEM (BF-TEM) in biological sciences is due, in part, to phase contrast being dominant in low atomic number soft matter samples ^11^. However, spatial resolution in BF-TEM decreases significantly beyond thicknesses greater than the elastic mean free path of electrons in water (*λ* ∼ 300 nm at 300 kV ^12^). In addition, as the sample thickness increases the flux in the elastically scattered wavefield decreases reducing the phase contrast even for zero loss filtered images. As most prokaryotic and eukaryotic cells are greater than 500 nm in thickness, conventional BF-TEM has limitations for high-resolution imaging of whole cell samples. Annular darkfield STEM (ADF-STEM) using incoherently scattered electron phase information, can provide superior spatial resolution as it is less affected by the effects of chromatic aberration ^13^, although at the expense of a reduced signal due to weak scattering at high angles from light element materials. The ADF-STEM geometry can also provide valuable complementary information on the elemental composition of biological samples using energy dispersive X-ray (EDX) spectroscopy which, for example, enables the localisation of biominerals with distinct elemental signatures in bacteria ^14^ and in tissue sections to distinguish between different heavy metal conjugated immuno-labels ^15^.

While ADF-STEM can improve image contrast over BF-TEM for thicker samples, the total thickness of the liquid cell in which the specimen is enclosed plays an equally significant role in limiting the achievable resolution and signal-to-noise ratios (SNR) of both LP-ADF-STEM imaging and EDX spectroscopy ^16^. This sample thickness limitation is compounded further by the need for samples to be enclosed in thin membrane windows to partition the sample from the microscope vacuum, e.g. each silicon nitride (Si_3_N_4_) window in dedicated TEM liquid holders is typically ∼50 nm thick, in addition to the liquid/sample thickness. Graphene liquid cells (GLC) can overcome some of the effects from the geometry of commercially available liquid holders, minimising the liquid encapsulation in a single atomic layer, therefore maximising the achievable imaging and spectral resolution from aqueous environments in LP-EM applications.

Selection of appropriate biological models are also important to achieving high resolution images of important biology using LP-EM. The *Deinococcus* genus is distinguished by an extraordinarily high tolerance to the effects of environmental hazards, such as arsenic ^17^, ionizing and UV irradiation ^18^, and desiccation ^19^. *D. radiodurans*, the type species of the genus, is a spherical, non-motile, didermal bacterium and is one of the most radiation-resistant of the *Deinococcus* genus, thirty times more resistant than the more commonly studied, *Escherichia coli* ^20^. While the biochemical mechanisms underlying this resistance are not yet fully understood, it is thought that this extreme radiation resistance evolved from its adaptation to dry environments ^21^ with dehydration being a more common physiological stressor than environmental radiation. Mechanisms for the unusual radiation resistance of *D*.*radiodurans* have been suggested to involve DNA damage resistance and repair pathways, as evidenced by the unusually large number of (>100) radiation-induced double-strand breaks (DSBs) that this species can withstand without lethality or mutagenesis. The proteins responsible for these resistance and repair pathways require protection from the oxidative environments created by irradiation and are therefore highly dependent on radical oxygen scavenger Mn^2+^ complexes ^22^ and antioxidant carotenoids such as deinoxanthin ^23^. These chemicals require a rich source of nutrients, to maintain effective concentrations. These resistance mechanisms, in turn, may allow *D*.*radiodurans* to tolerate electron irradiation for longer, mitigating the effects of electron beam damage ^24^. *D*.*radiodurans* is therefore an ideal candidate for ultrastructural study using LP-EM techniques.

In this work, we demonstrate the capabilities of graphene encapsulation and STEM-EDX imaging in studies of biological specimens in the liquid phase. We use these to provide insights into *D*.*radiodurans* ultrastructure using graphene encapsulating environments, localising manganese ions in phosphate-containing storage granules within the bacterium and demonstrate the ultrastructural response of *D*.*radiodurans* to nutrient restriction and desiccation *in vivo*.

## Results

### Graphene liquid cell assembly and morphology of *D*.*radiodurans*

Using the graphene encapsulation method developed previously ^25,26^, it is possible to confine biological samples within graphene encapsulated aqueous environments in standard Au-flat and Au Quantifoil grids (Fig.1A-E). A single-layer graphene grid was submerged in a container with a biological buffer of interest and the sample was pipetted over it. The second layer of graphene was then carefully lowered onto the grid by removing excess buffer from the container and allowed to dry. The encapsulated bacteria were immediately inspected using brightfield light microscopy (LM). The graphene encapsulation appears to be tolerated by *D*.*radiodurans* as shown by the integrity of the membrane under visual inspection in LM and EM (Fig.S1A-B). Thus, we calculated the approximate value of the sustained pressure on a *D*.*radiodurans* cell, using the Laplace pressure ^29^, yielding an approximate pressure of 0.5 atmosphere (∼50 kPa) (Fig. S2, S3). A more detailed methodology includes different pressure contributions such as Van der Waals and elastic components ^30^, which yields a pressure around 1.6 atmospheres (∼162 kPa). This result is in good agreement with both the measured intactness of the membrane ^31^ and the calculated pressure for samples as thick as bacteria or cells (1 μm) using different parameters ^29^. In the case that a direct contact between the pristine graphene and the hydrated membrane of *D*.*radiodurans* is considered, the estimated pressure is several orders of magnitude lower. See analysis in supplementary information for more details. In all lysogeny broth (LB) cultures, *D*.*radiodurans* cells typically formed tightly packed tetrads, with a high degree of asymmetric division and poorly formed central septa in early exponential growth phases and predominantly octads with thick central septa in stationary phase cultures (Fig. S1C-F). In low nutrient media preparations, the cells primarily formed single cocci or diads, as observed previously in studies of *D*.*radiodurans* nutrient-dependent pleomorphisms ^27,28^.

**Fig.1.**
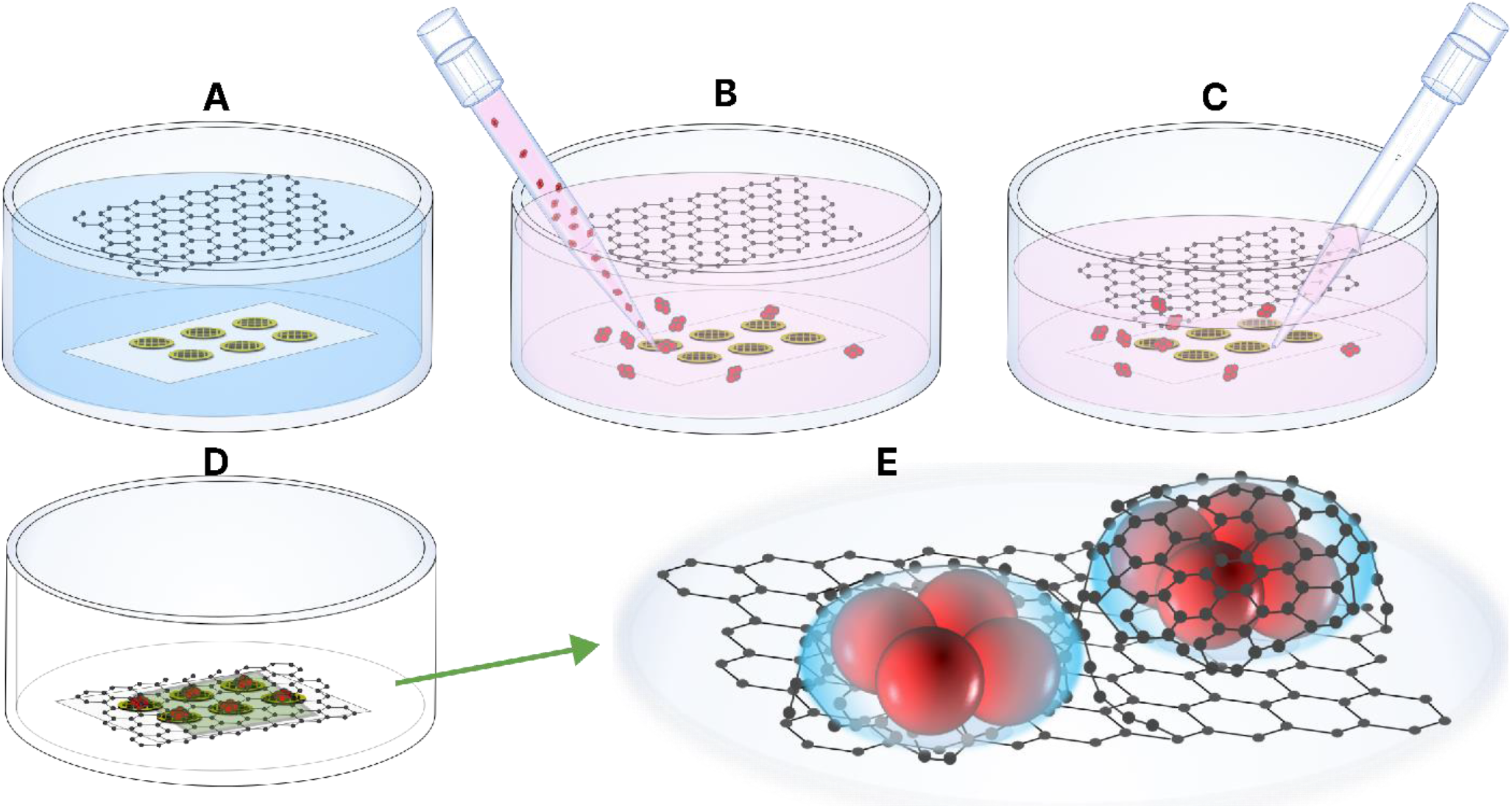
Schematic of graphene blister assembly and LM/EM analysis of encapsulated *D*.*radiodurans*. **A)** Freshly prepared single-layer graphene grids placed in water on filter paper, with the second layer of graphene suspended above. **B)** After exchanging water with buffer (HEPES), the sample is added directly to the top of the graphene grids. **C)** Solution is removed slowly to ensure the graphene layer remains intact over the grids. **D)** Solution is completely removed, and grids allowed to air dry. **E)** Expanded view of panel **(D)** of graphene blisters (GB) with encapsulated *D*.*radiodurans*.

### *D*.*radiodurans* growth stages, ultrastructure, and elemental distribution

Imaging whole bacteria provides cellular context to subcellular features, allowing rapid screening of multiple bacteria at different stages of growth and capturing rare cellular events which may be obscured by sample thickness or removed by thinning in conventional cryo-EM studies ^32^.

Bacteria were encapsulated at different phases of the bacterial growth curve; stationary phase bacteria were often found in groups of octads, with multiple high contrast granules present. The periplasmic space between the outer and inner membrane (OM and IM, respectively) was ∼90 nm (Fig.2A-B), with a peptidoglycan (PG) component ∼35 nm thick and an interstitial layer (IL) ∼55 nm thick. This agrees with previously reported descriptions of a dense diffuse periplasmic layer ^33^ and cell envelope thickness measurements from frozen vitrified sections ^34^ of *D*.*radiodurans*. The surface layer (S-layer), typically reported as a 20 nm thick layer ^34^, was not clearly visible. However, a high contrast layer (HCL), ∼55 nm thick, likely containing the S-layer and carbohydrate coat ^35–38^, was observed (Fig.2B).

**Fig.2.**
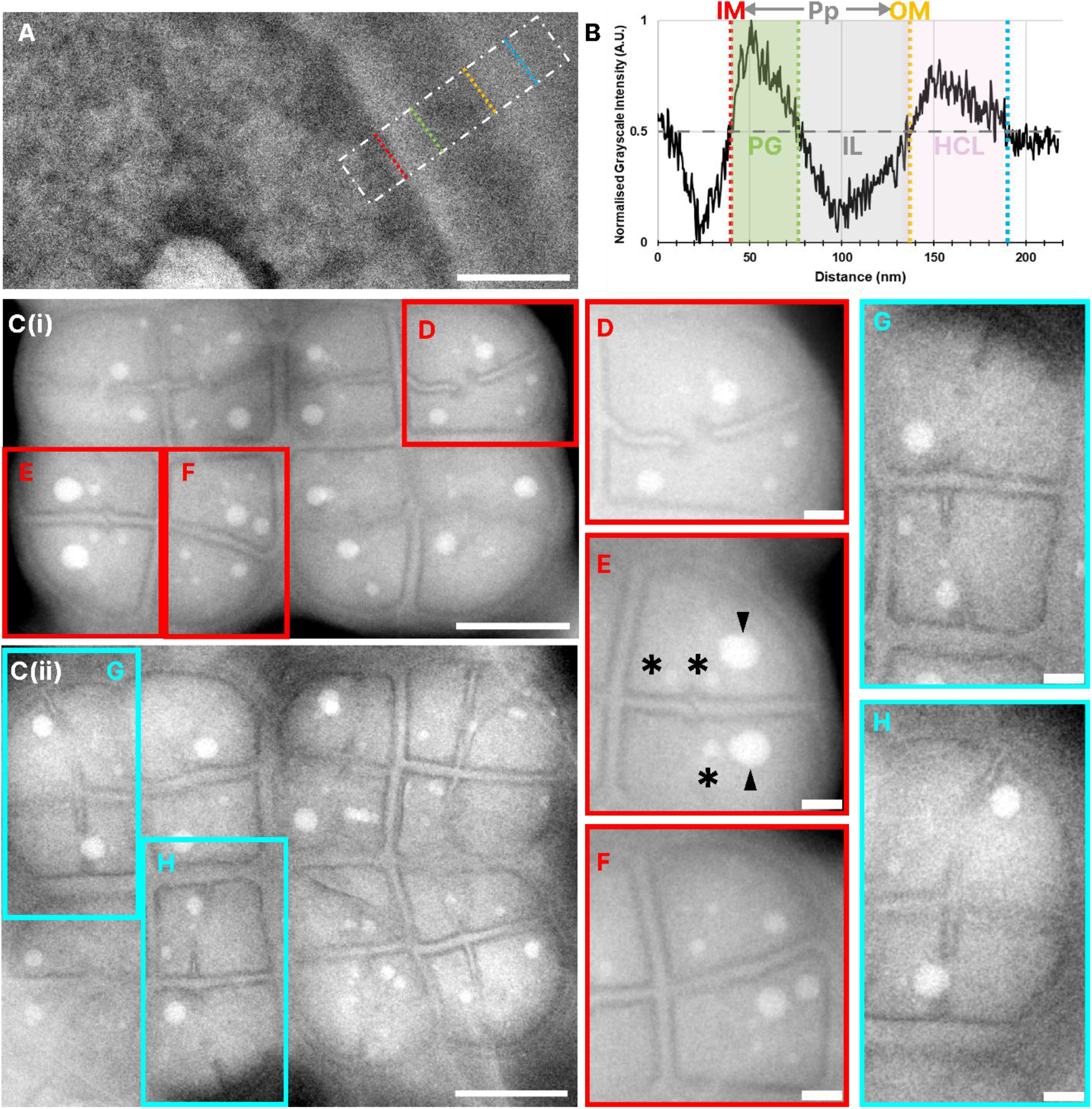
*D*.*radiodurans* growth stages and membrane composition in graphene encapsulated environments. **A)** Bandpass (BP) filtered LP-ADF-STEM section of encapsulated late exponential phase (OD_600_: 0.65) *D*.*radiodurans* cell envelope, with the density profile region of interest (ROI) highlighted (TCF: 37,500 e^-^/nm^2^). **B)** Corresponding density profile of cell envelope components reveal a periplasmic space (Pp) between the inner and outer membrane, a peptidoglycan layer (PG) followed by an interstitial layer (IL) and a final high contrast layer (HCL). **C i-ii)** BP filtered LP-ADF-STEM images of encapsulated stationary phase *D*.*radiodurans* octads at various stages of division from the same sample (TCF: 93 e^-^/nm^2^). **D-H):** Expanded view of single cocci from octad ROIs showing septum progression stages. **D)** Characteristic septal aversions visible. **E)** Septa meeting side on, in the cell centre. Asterisk (*): Small storage granules; Arrowheads: Phosphate storage granules. **F)** Two cocci separated by a completed septum. **G)** New septa begin to emerge perpendicular to the central septum. **H)** Later stage of tetrad formation. Scale bar: A: 100 nm; C i-ii: 1 μm, D-H: 250 nm.

The characteristic stages of septal growth and cell division in *D*.*radiodurans* were observed in encapsulated bacteria (Fig.2Ci-ii). Septal growth begins perpendicular to the central septum of the tetrad, dividing a single coccus into two cocci with characteristic septal aversion, as previously reported in conventional EM studies ^39^ (Fig.2D, E). Large phosphate granules were evident in the majority of *D*.*radiodurans* cocci, with smaller storage granules ranging from 40 – 150 nm in size, potentially used for carbohydrate storage often visible ^40^, as highlighted in Fig.2E. In early exponential cultures, the phosphate granules appeared smaller, were difficult to differentiate from carbohydrate granules and were also more numerous (Fig. S1A). Merging of the septa leads to complete cellular division into two cocci (Fig.2F). Further division into four cocci begins with the formation of septa in the centre of and perpendicular to the previous septum and on opposite sides of the cell walls (Fig.2G). The septa advance from the central and peripheral edges toward the centre of the coccus with a diminishing slit formed between the advancing septal edges (Fig.2H), as observed previously by optical microscopy ^41^ and conventional EM^39^.

Several asymmetric divisions of cocci were also observed (Fig. S4A), along with s-shaped septal growths from the centre of the cell (Fig. S4 B, C) and multiple diagonally orientated septa at different stages of division (Fig. S4D, E). It was possible to image these structures at fluences as low as ∼0.5 e^-^/nm^2^ (Fig. S5). A fluence of 0.5 e^-^/nm^2^ is an order of magnitude lower than the fluence threshold for maintaining transcriptionally active *E*.*coli*, suggesting live cell imaging of these structures may therefore be possible in the near future ^42^. Note, while the terms electron dose and electron fluence are often used interchangeably in the field of electron microscopy, electron fluence is defined as the number of impacted electrons on a sample per unit area, i.e. 100 e^-^/nm^2^ = 1 e^-^/Å^2^, and electron dose is strictly defined as the energy absorbed per unit area, expressed in units of Grays. Therefore, in all subsequent images we quote the total cumulative fluence (TCF) in SI units of e^-^/nm^2^, which accounts for all electron beam irradiation up to and including the image shown.

At higher magnifications, some ultrastructural features were visible within the encapsulated cells (Fig.3A; Fig. S6A-D), including protein (P), DNA, peptidoglycan (PG), septal edges (S) and storage granules (SG). Protein was dispersed throughout the cell and power spectra (PS) calculated from image areas containing protein (Fig.3B, red ROI), show enhanced frequencies corresponding to a spacing of 24 ± 8 nm consistent with bacterial ribosomes, not observed in the PS calculated form the peptidoglycan region, blue ROI (Fig. 3B, Inset; Fig. S6 E-H).

**Fig.3.**
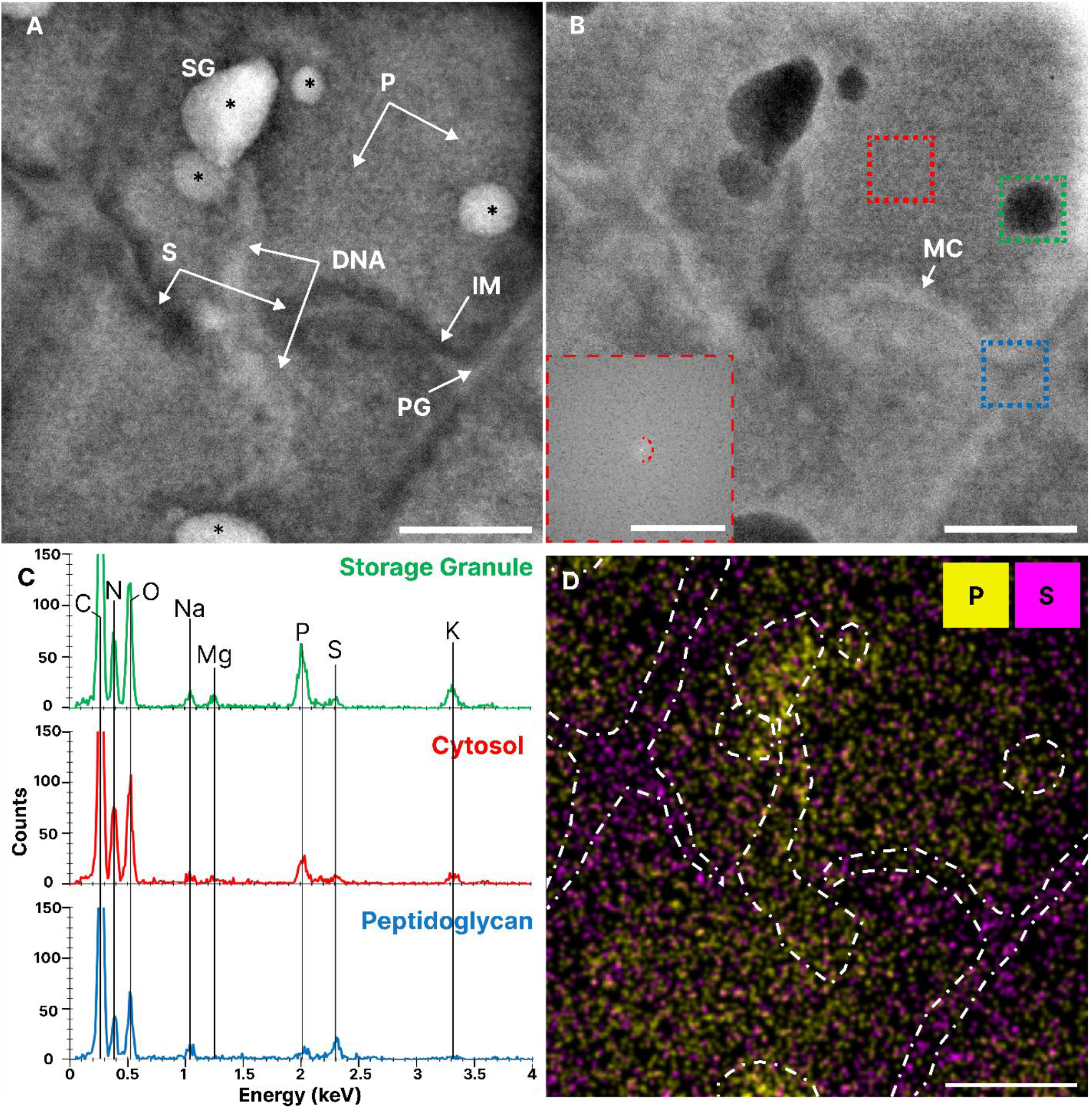
Ultrastructural detail and elemental distribution in *D*.*radiodurans* using LP-ADF-STEM and EDX. **A)** BP filtered LP-ADF-STEM and **B)** LP-BF-STEM image of encapsulated bacteria internal structure (TCF: 11,800 e^-^/nm^2^), **Inset:** PS of cytosolic (red) ROI, with spatial frequency at (15 nm)^-1^ marked (dashed red). **C)** EDX spectra of ROIs in **(B). D)** Gaussian filtered and contrast normalized phosphorous and sulphur EDX map of **(A, B)** (TCF: 50,300 e^-^/nm^2^). Scale bar for A, B,D: 250 nm; PS: (2 nm^-1^).

EDX analysis of the IM and PG indicates a relatively high abundance of sulphur. We propose that this periplasmic layer may act as a reservoir for sulphur-containing amino acids and low molecular weight thiols, such as cysteine/methionine and bacillithiol, respectively, which are integral to the cells redox homoeostasis and antioxidant defence system ^43–45^. Most storage granules imaged, were enriched in phosphorous, oxygen and potassium (Fig.3C, Fig. S7). Other counterions including, magnesium were also observed to correlate spatially with phosphate granules although the EDX SNR was poor due to the relatively low fluence used for acquisition. While calcium counterions were also observed in polyphosphates with *D*.*radiodurans*, these were relatively rare events and usually only in single polyphosphate granules within desiccated individuals. Sulphur was found to correlate with the cell membrane, along with sodium and chlorine (Fig. 3D, 4).

**Fig.4.**
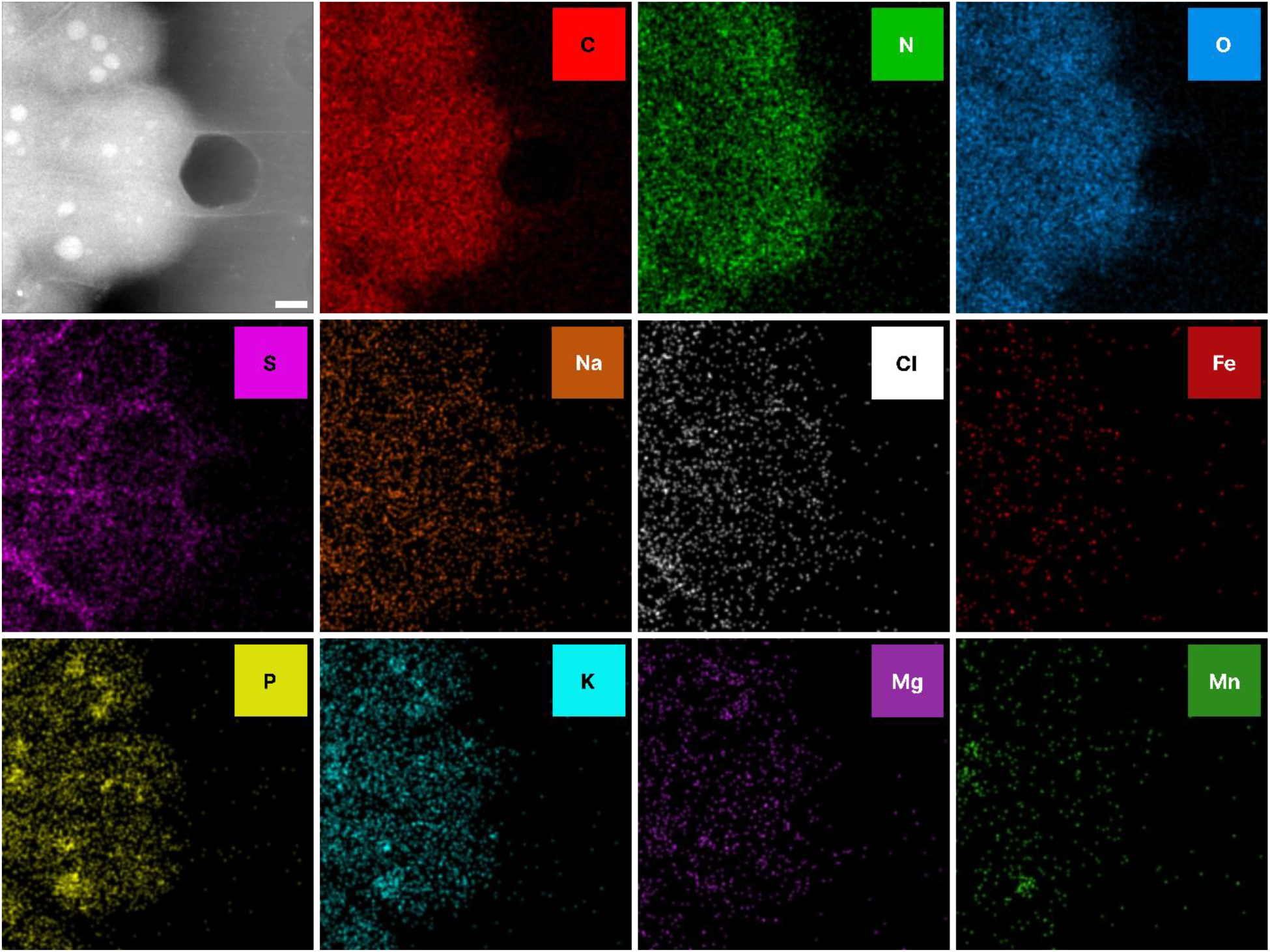
Manganese accumulates in bacterial phosphate granules. Summed LP-ADF-STEM image of bacteria incubated in MnCl_2_ (Average TCF: 2,979 e^-^/nm^2^). **Top Row:** Intracellular elements; **Middle row:** Cell membrane associated elements; **Bottom Row:** Polyphosphate associated elements. Scale bar: 500 nm

Spherical, low contrast particles were also observed, roughly 30 nm in size, associated with the IM on the cytoplasmic side and are likely to be similar structures to those previously reported as macromolecular complexes (MC) ^34^, thought to be Dps2 clusters for iron sequestration ^22^ (Fig.3B). However, there was no significant iron signal found anywhere within the cell in the EDX maps (Fig. 4). This is possibly due to iron being dispersed throughout the cell rather than being concentrated in any specific area making the signal extremely weak in an already relatively low signal map (Fig. S7B).

### Manganese accumulates in bacterial phosphate granules

Manganese is implicated in *D*.*radiodurans* radiation resistance and appear to localise with phosphate ions ^22^ suggesting that manganese acts as a counterion to cellular phosphate. Using EDX analysis, at biologically relevant fluences i.e. <3,000 e^-^/nm^2 46^, we confirm that manganese localizes to phosphate granules following MnCl_2_ supplementation of bacterial media (Fig. 4; Fig. S8A). Interestingly, manganese is not distributed uniformly across all polyphosphates within the cell, suggesting distinct biological roles for the different polyphosphates-cation identities. However, we were unable to observe accumulations of naturally occurring manganese-containing polyphosphates in *D*.*radiodurans* at the fluences used to record the maps (Fig. S8 B, C). These manganese-containing phosphate granules may act as a store of reactive oxygen species-scavenging manganese cations for release during oxidative stress.

### Vacuum desiccation in *D*.*radiodurans* leads to separation of cell membrane, a marker for liquid encapsulation

To assess the effect of vacuum desiccation on *D*.*radiodurans*, a sample exponentially growing bacteria were placed on a graphene grid as previously mentioned, however after a short period (<5 mins) of drying the grid was placed directly into the microscope. Bacteria exposed to the vacuum detached from their membranes causing large cracks to form (Fig.5A). These characteristic cracks, act as an internal standard for verification of encapsulation, making selection of regions of interest (ROI) straightforward even at relatively low fluences (<1 e^-^/nm^2^). This reduces the requisite pre-acquisition fluence budget ^47^, thus allowing more fluence to be used for single frame high resolution imaging or multi-frame, dynamic time course experiments.

**Fig.5.**
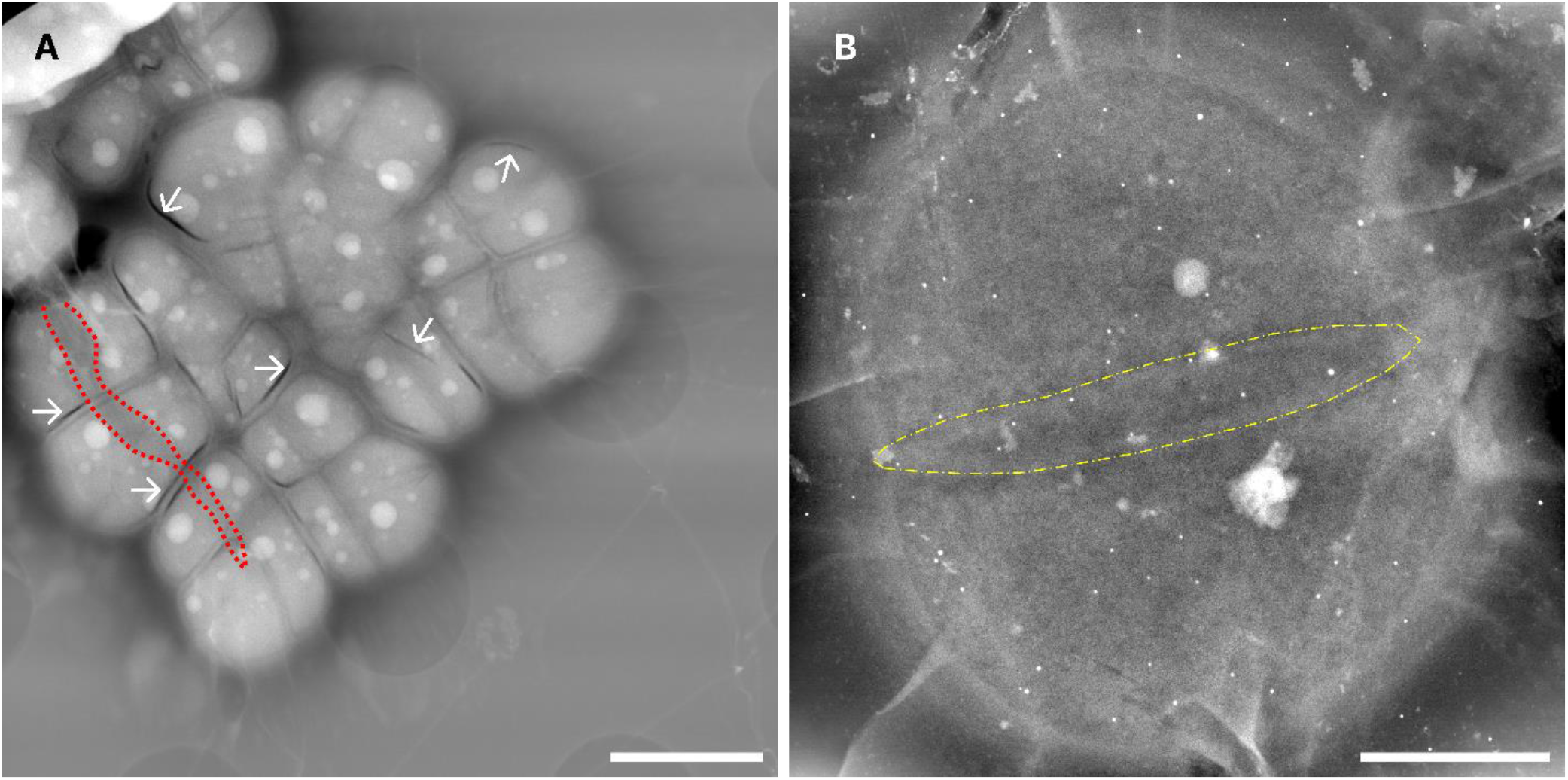
Effect of different environmental stressors on *D*.*radiodurans* ultrastructure. **A)** LP-ADF-STEM image of vacuum desiccated bacteria in a torn graphene liquid cell (TCF: 93 e^-^/nm^2^); Dashed red line: Tear in graphene, White arrows: Examples of desiccation-related membrane separation. **B)** LP-ADF-STEM image of encapsulated diad, grown in 10% LB media (TCF: 2,334 e^-^/nm^2^); Dashed yellow line: Ring shaped membrane between individuals. Scale bar for A, C: 500 nm; B: 2 μm.

### Nutrient restriction leads to enrichment of different *D*.*radiodurans* pleomorphs

As *D*.*radiourans* demonstrates nutrient-induced pleomorphisms ^27^, we decided to grow *D*.*radiodurans* in nutrient-restricted conditions (10% v/v LB media) in an effort to explore the different ultrastructure of these pleomorphs and to create smaller cell volumes to potentially enable three-dimensional data acquisition using LP-STEM tomography. It was readily apparent using light microscopy that nutrient-restriction dramatically reduces the number of tetrads and increases the number of monad and diads in solution. This was confirmed in the electron microscope, as most bacteria present were diad forms. While the diads did appear to have a less dense cytoplasm than tetrads there was little difference in the ultrastructural features, with the primary difference being a more spherical shape rather than ellipsoidal. Interestingly, in some individuals it was clear that the peptidoglycan wall between two individuals was in a ring shape rather than a singular partition suggesting that there may be significant intercell communication in these diads at this stage of growth (Fig.5B). We also observed diads which do not display this ring-shaped aperture and instead had a typical peptidoglycan partition, these appeared more ellipsoidal, suggesting that there may be a need for this type of aperture at this specific stage of cell growth. A similar decrease in the ellipticity of the cells at specific growth phases has been observed previously by high resolution light microscopy ^41^.

## Discussion

The LP-STEM studies presented here demonstrate an effective technique for imaging whole biological specimens and their internal ultrastructure in native aqueous environments, without the need for heavy-metal staining, nanoparticle tagging, cryogenic/plastic preservation requiring specialized equipment and leading to potential artefacts. The graphene liquid cells used, prepared from relatively simple materials, provide high contrast for LP-EM analysis. In combination with STEM-EDX techniques, they can provide greater internal information than other native preparation approaches, such as conventional whole-cell cryo-transmission electron microscopy/tomography techniques where samples contain layers of vitreous ice.

As the graphene liquid cells are prepared on EM grids, they can fit in most standard EM holders. Unlike commercially available silicon nitride chip liquid cells, the nature of GLCs means each graphene blister is an independent liquid cell, keeping potential hydrolysis products from the electron beam localised to a single blister and preventing the build-up and diffusion of free radicals casing damage in adjacent cells. The isolated nature of graphene blisters also means that in the event of a failure of a single liquid cell, there is no significant release of liquid inside the microscope column. This contrasts with conventional silicon nitride chip liquid cell applications where failure of the chip may lead to the release of larger volumes of liquid and gases in the column and where great care is required to ensure the liquid cell remains sealed. However, the most significant advance in imaging quality is the electron transparency of the graphene and thinner liquid volume.

There are several variables that we found to significantly affect sample quality and sensitivity and image SNR. For example, we found that while gold Au-flat grids produced a tighter encapsulation and more reproducible number of blisters, gold Quantifoil grids gave images with better SNR over larger fields-of-view. Additionally, buffer composition is a significant factor; the use of phosphate buffered saline led to substantial amounts of salt precipitates on the grid obscuring the field-of-view. We therefore used a simpler zwitterionic buffer, HEPES, for our experiments. Degassing before encapsulation was also essential to limit the formation of gas bubbles in the buffer under the electron beam.

Our study emphasises the importance of selecting suitable model organisms for study in LP-EM. *D*.*radiodurans* has proven to be an extremely valuable test sample for exploring LP-EM/EDX capabilities due to its radiation resistance, elementally heterogenous nature and round coccoid shape, which lends itself readily to study in GLCs. Previously, it was shown that in transcriptionally active *E*.*coli*, 50% of the population received a lethal fluence (LF_50_) at ∼10 e^-^/nm^2^, in commercially available silicon nitride chips ^42^. However, *D*.*radiodurans* is known to tolerate fluences at least tenfold greater in magnitude than *E*.*coli*, suggesting an LF_50_ of *D*.*radiodurans* under the same circumstances of approximately 100 e^-^/nm^2^. This fluence limit is further increased by the 2-7x protection factor afforded by graphene encapsulation ^48–50^, potentially leading the way for low fluence live-cell imaging in LP-STEM. Our observations of proteins at ∼24 nm and other biomolecules within these bacterial cells suggest that even at cumulative fluences significantly higher than the putative LF_50_ of *D*.*radiodurans*, it may still be more favorable to monitor protein dynamics and interactions *in situ* than in purified protein solutions, with beam-related damage potentially mitigated by the native reactive oxygen defense system of *D*.*radiodurans. D*.*radiodurans* response to desiccation by creating cracks in their membrane from rapid depressurization in the microscope vacuum, also demonstrate their utility in LPEM analysis, by acting as an internal marker for aqueous environments within the electron microscope.

We have also demonstrated the use of LP-STEM-EDX applications to build an elemental understanding of the bacterial cell at the ultrastructural level. LP-STEM-EDX has enabled us to demonstrate that Mn ions localize to polyphosphate granules *in situ*, without the need for denaturants and fixatives. We have also mapped significant concentrations of sulphur residing in the periplasm, suggesting a potential mechanism for oxidative homeostasis in *D*.*radiodurans*. This periplasmic sulphur may help manage redox homeostasis within *D*.*radiodurans*, being released into the cytoplasm on irradiation to help inhibit oxidative damage induced by free radicals and protect biomolecules critical to cellular survival. These LP-STEM-EDX studies demonstrate the potential to use both native elemental distributions and non-native elemental probes to localise biomolecules and structures of interest within a cell.

Finally, this study is a proof of principal showing how LP-STEM-EDX enables the analysis of large-scale cellular events, such as cell division, under relatively low fluence conditions (<100 e^-^/nm^2^) and can resolve even finer ultrastructural details, such as proteins and nucleic acids albeit at a higher fluence. The techniques described herein illustrate the potential of LP-EM to explore fundamental biological questions *in situ*. Electron fluence, sample thickness and the fundamental characteristics of graphene, such as its grain size, may however limit its broader application to larger cells and tissues. Ultimately, LP-EM techniques combined with emerging low-fluence imaging applications such as ptychography ^51,52^ and compressive sensing ^53^ mean that imaging cellular dynamics *in vivo* at sub-nanometre resolutions in an electron microscope may soon be achievable.

## Materials and Methods

### Growth of *D. radiodurans*

5 ml of Lysogeny Broth (LB) was inoculated with *D. radiodurans* (Strain R_1_; DSMZ 20539) frozen culture and incubated at 30 °C for 24hrs with shaking. 500 μl of the resulting culture was added to 5 ml of fresh LB [1% w/v Glucose] and after this reached early exponential phase (OD_600_: ∼0.3) was further incubated and then collected at different time points: 15 min (early exponential phase, A600 nm = 0.35), 2 h (mid-exponential phase, A600 nm = 0.65) and 20 h (stationary phase, A600 nm = 1.65). For the Mn accumulation experiment, early exponential phase bacteria were incubated with sterile-filtered MnCl_2_ [0.1 or 1 mM] for 4hrs, then spun down and encapsulated. A control growth with water was performed and followed simultaneously. For the restrictive media experiment, 500 μl of stationary phase bacteria were added to 5ml of fresh 10% LB media in ddH_2_O [1% w/v Glucose] and incubated overnight. 2ml of the resulting samples were then spun down at 500g for the LB preparations and 2000g for the restrictive preparations for 5 mins, resuspended in 1 ml of nitrogen degassed 4-(2-hydroxyethyl)-1-piperazineethanesulfonic acid (HEPES) buffer [25 mM] and encapsulated in graphene for LP-STEM/EDX imaging.

### Brightfield light microscopy imaging of DAPI-stained *D*.*radiodurans*

Imaging was carried out with an upright brightfield light microscope, Eclipse Ni (Nikon, USA) using 10x and 40x air immersion objectives.

### Graphene liquid cell encapsulation

The hydrated graphene enclosed environments were obtained by first creating a graphene TEM grid by transferring graphene to a TEM grid using cellulose acetate butyrate ^54^ followed by a dry-cleaning procedure ^55^ at 310 °C overnight. This is the basis of the liquid cell. The top graphene layer for encapsulation was fabricated using a polymer-free transfer method ^56^, which provides a flat top surface to the hydrated environment which then adapts to the morphology of the target. The main steps are described in detail in refs ^57^ and ^26^ to prepare the base TEM grid and the creation of the encapsulated environments respectively ^58^.

The commercial based TEM grids used throughout the different experiments to create the hydrated encapsulated environments were Au-Flat (2/2) 300 Mesh, 45nm Thick from Protochips and Au Quantifoil R1.2/1.3 300 mesh from Quantifoil.

### Liquid phase STEM imaging

All STEM imaging was performed on a 300kV GRAND-ARM 2 (JEOL) with a spherical aberration corrector, with an illumination convergence semi-angle (α) of 15.95 mrad and annular dark-field (ADF) [Inner collection semi-angle (β^in^: 37.1 mrad)/Outer collection semi-angle (β^out^: 83 mrad), measurement limited by the differential pumping aperture and bright-field (BF) detectors (β_BF_: 20.8 mrad). The microscope was aligned using the standard gold waffle thin film. Images of aqueous bacteria were typically recorded at 20 kX, 100 kX and 200 kX magnifications (pixel size: 5, 1 and 0.5 nm respectively), 2048 x 2048 frame size and 20 μs dwell time.

### Liquid phase EDX imaging

Energy-dispersive X-ray spectroscopy was performed in STEM mode on a 300kV GRAND-ARM 2 (JEOL) on aqueous bacterial samples at 20kX, 100 kX and 200 kX magnifications (pixel size: 20, 4 and 2 nm) respectively), 512 x 512 frame size and 20 μs dwell times, using dual liquid N_2_ cooled Si (Li) JEOL detectors with a combined solid angle of 1.41 sr.

### STEM/EDX Image preprocessing

To enhance high-resolution details and to remove background noise, a Gaussian bandpass filter was applied to each micrograph (High Pass :300 px/Low Pass:2 px), using the ImageJ Bandpass filter plugin. Horizontal STEM scanning artefacts were smoothed using the “Suppress Stripes” function at 5% in the ImageJ Bandpass filter plugin. All images were then normalised with 0.35% pixel saturation, using the “Contrast Enhance” function in ImageJ.

EDX maps were produced in JEOL Analysis Station Software and processed in Hyperspy ^59^. Maps were binned by 2 in x-y and Gaussian filtered (*σ* = 2 px) to remove noise, using the “Gaussian Blur” function in ImageJ. All maps were then contrast normalised with 0% pixel saturation, using the “Contrast Enhance” function in ImageJ. All maps were generated using the respective elemental Kα lines.

### Fluence History Tracking with Axon Studio

The Axon Studio software from Protochips Inc., was used in conjunction with Axon Dose to accurately track the fluence history of each imaged area, providing an accurate readout of specimen total cumulative fluence.

## Acknowledgments

We would like to thank Tobias Starborg, Yusuf Mohammed for help with equipment and valuable discussions and Chen Huang for help setting up the electron microscope for EDX.

## Author contributions

B.C. conceptualization; B.C. and A.P.T. investigation; B.C., A.P.T., E.L., B.G., J.K. and A.I.K methodology; B.C. and A.P.T. formal analysis; B.G. software, A.P.T. resources; B.C. and A.P.T. visualisation, B.C. and A.P.T validation, B.C., A.P.T., J.K. and A.I.K. writing – original draft, B.C., A.P.T., B.G., E.L., B.G., J.K. and A.I.K. writing – review & editing, B.C., J.K. and A.I.K project administration.

## Data and materials availability

The data that support this study are available from the corresponding authors upon request. The raw data, LM, ADF-/BF-STEM, EDX and TEM images from this study have been deposited in the Zenodo database [https://doi.org/10.5281/zenodo.10625794]

## Supplementary Materials

### <u>Analysis of sustained pressures inside graphene encapsulated</u> *D*.*radiodurans*

We have measured the average size of the *D*.*radiodurans*. Based on the measurements from 13 bacteria from ADF-STEM images, *D*.*radiodurans* has an ellipsoidal geometry. The average lengths of the long axis and short axis are 2.91 ± 0.23 μm and 2.09 ± 0.13 μm, respectively. These values include both random and systematic errors. The standard deviations of the lengths of the long axis and short axis are 0.57 and 0.32 μm, respectively (Fig.S2).

The pressure inside a graphene liquid cell can be estimated by using the pressure of Laplace^29^:

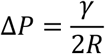

Where *ΔP* is the Laplace pressure gradient, γ the interfacial energy of graphene−water and *R* is the radius of the encapsulated *D*.*radiodurans*. The error associated with the calculated value is:

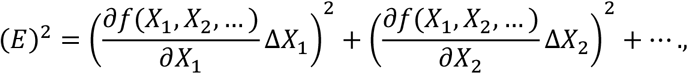

Thus:

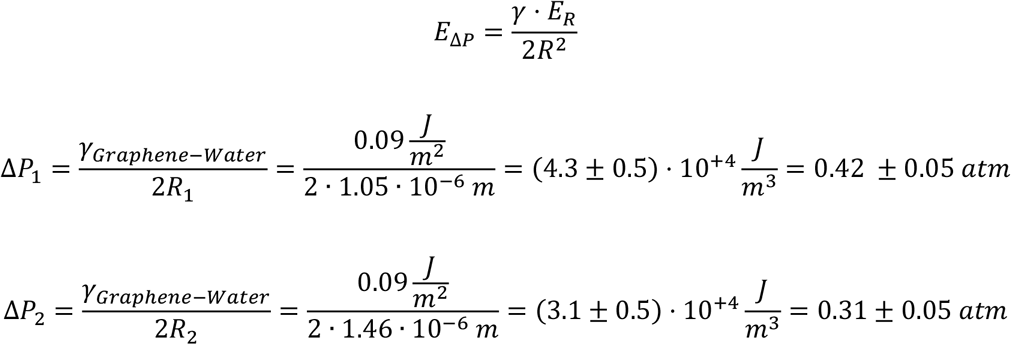

To calculate the pressure exerted by graphene directly, without intermediate water, on a hydrated *D*.*radiodurans*, some approximations can be applied. The first is that the membrane of a *D. radiodurans* is chemically similar to that of *E*.*coli*, and hence the surface tension can be estimated. Therefore, we can use the measured interaction force between a single flake of pristine graphene and the cell wall of a living *E*.*coli*: 38.2 ± 16.4 pN. Then, using the following equation ^31^ we can approximate the interfacial energy:

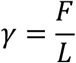

Where *F* is the adhesive force and *L* is length over which that force is distributed. In the case of *radiodurans* we can simplify its geometry to a circumference, where *d* is the diameter of the *radiodurans* which value is in the range of length of the long and short axis previously measured (2.1-2.9 μm):

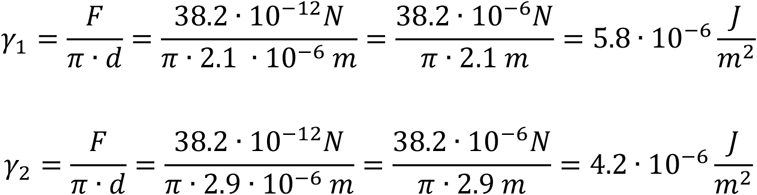

The error associated can be calculated using the same equation for error previously displayed, hence:

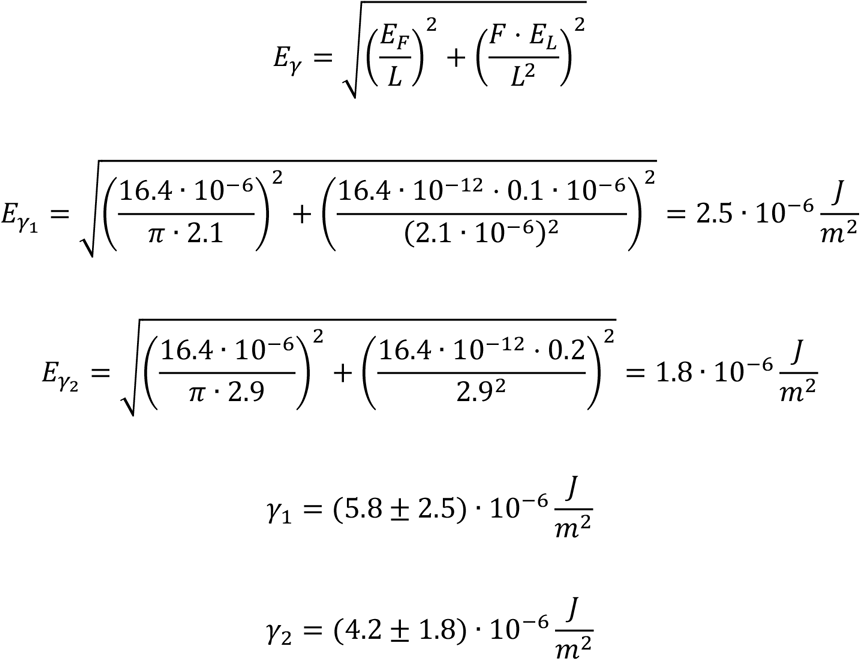

We can estimate the pressure within the encapsulated region from the surface tension exerted by pristine graphene on the membrane surface. This estimation is based on the Laplace pressure equation. The error in this estimation can be calculated using the following equation:

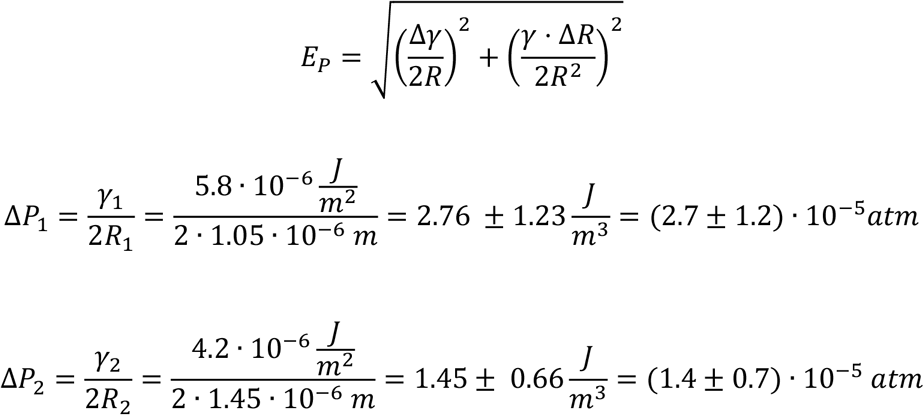

In addition, this pressure can be compared with the pressure calculated using a more detailed analysis which includes other pressure components such as Van der Waals and elastic, as performed by Khestanova *et al*^30^ in the following equation:

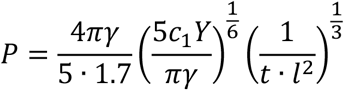

Where *γ* is the total adhesion energy (*γ* = *γ*_*Graphene*−*Graphene*_ − *γ*_*Graphene*−*Water*_, where *γ*_*Graphene*−*Graphene*_ and *γ*_*Graphene*−*Water*_ are the adhesion energy between two graphene layers and graphene and water respectively), *Y* is the Young’s modulus and *t* and *l* are the height and the radius of the encapsulated area, respectively. Substituting the variables by the associated values we reach almost twice atmospheric pressure:

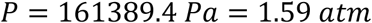

This equation can also be used to estimate the total pressure of other encapsulated biological structures, such as viruses and proteins, which are of utmost importance in future experiments (Fig. S3).

---

**Fig. S1.**
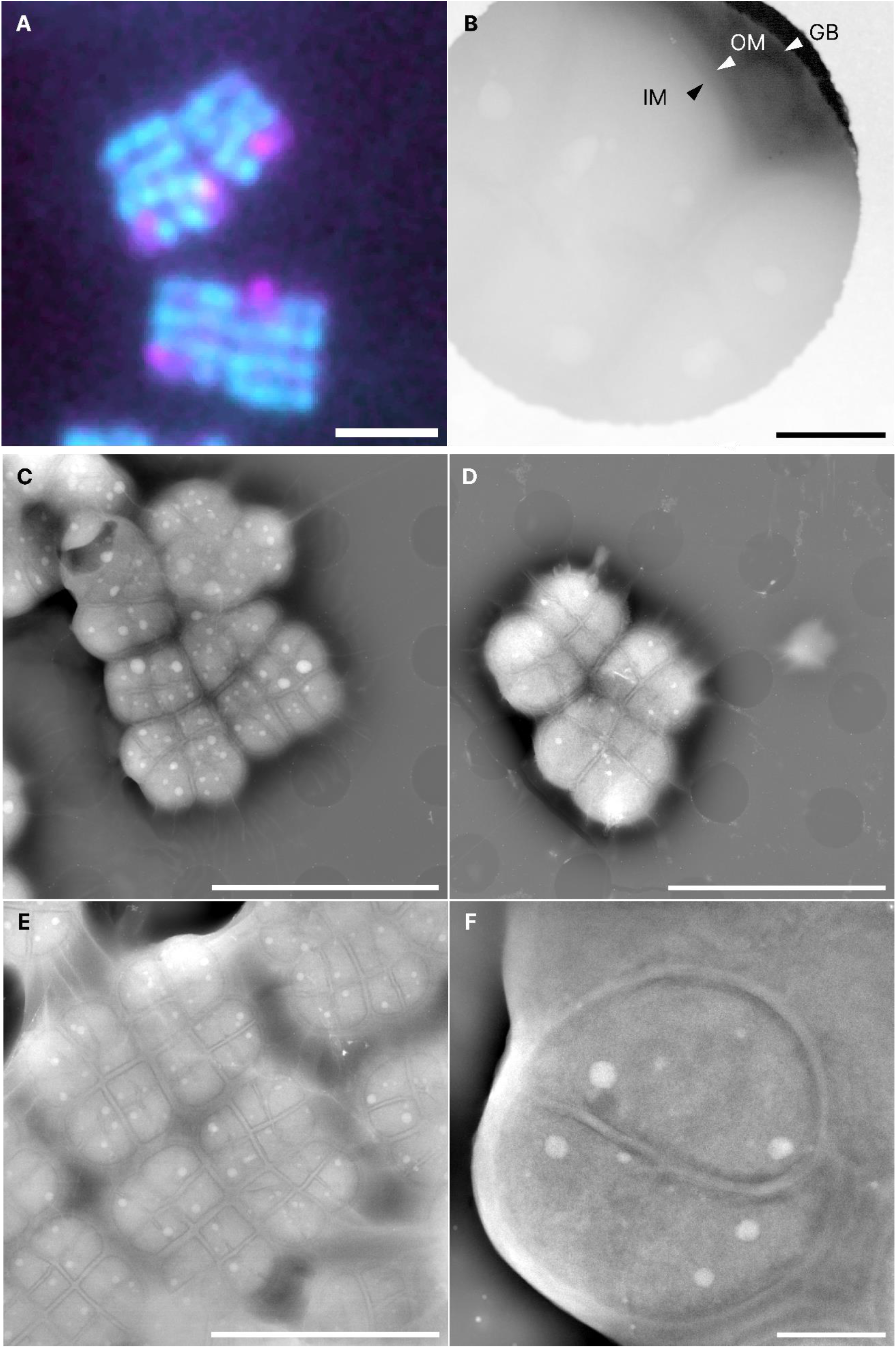
*D*.*radiodurans* morphology and growth stages in graphene encapsulated environments. **A)** Brightfield light microscope image of DAPI-stained *D*.*radiodurans* grown in LB broth, indicating nucleoid (cyan) and phosphate granules (magenta). **B)** LP-ADF-LPSTEM image (log grayscale) of encapsulated bacteria at 100kX, with both inner membrane (IM) and outer membrane (OM), along with graphene blister edge (GB) indicated. LP-ADF-STEM image of encapsulated early exponential phase (OD_600_: 0.35) **(C)**, late exponential phase (OD_600_: 0.65) **(D)** and late stationary phase (OD_600_:1.65) **(E)** *D*.*radiodurans*. (**F)** LP-ADF-STEM image of encapsulated *D*.*radiodurans* diad from the late stationary phase sample. (TCF: 93 e^-^/nm^2^) Scale bar: A,C-E: 5 μm. B,F: 500 nm.

**Fig. S2.**
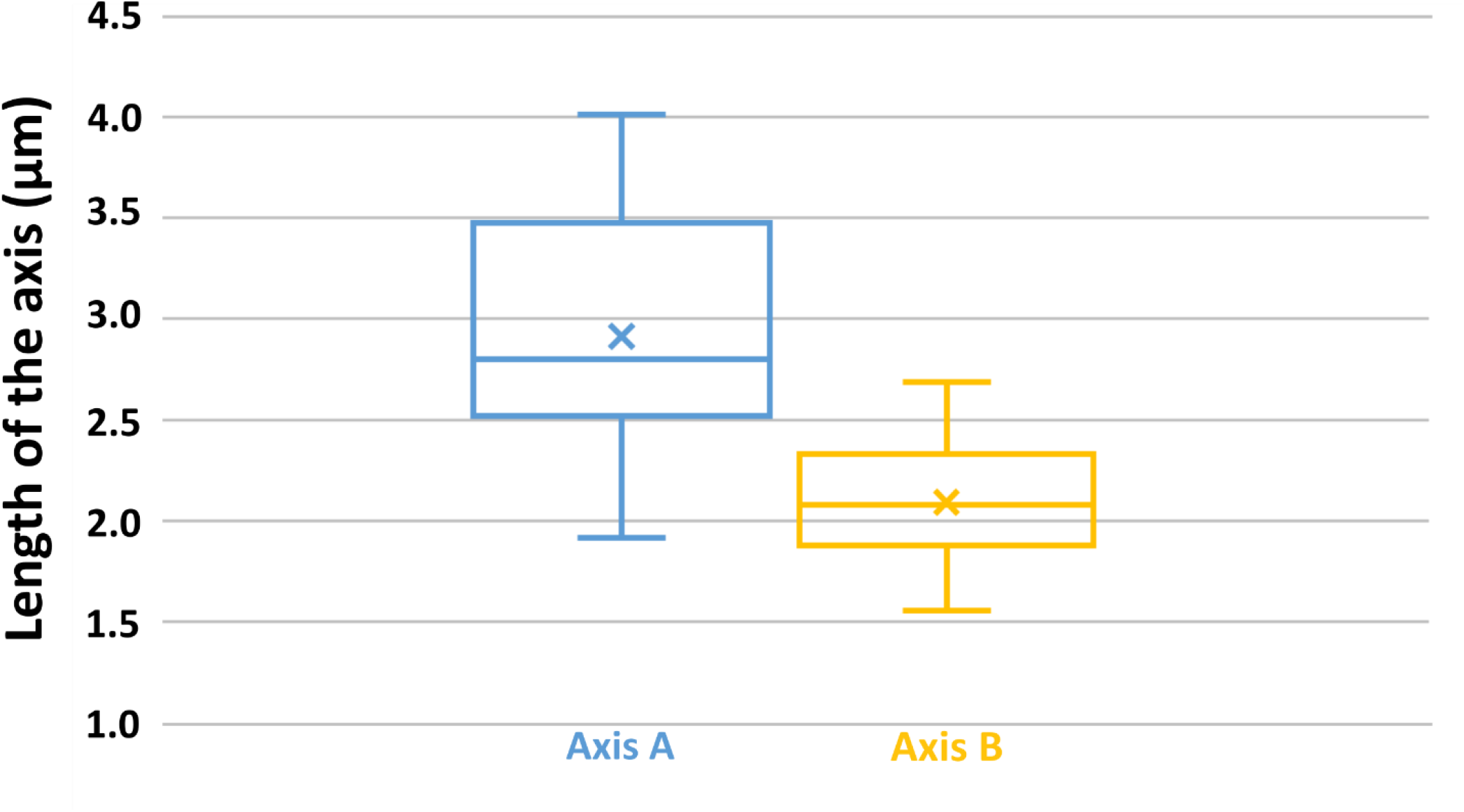
Statistical analysis of *D*.*radiodurans* size distribution. The centre line indicates the median values; × indicates the mean.

**Fig. S3.**
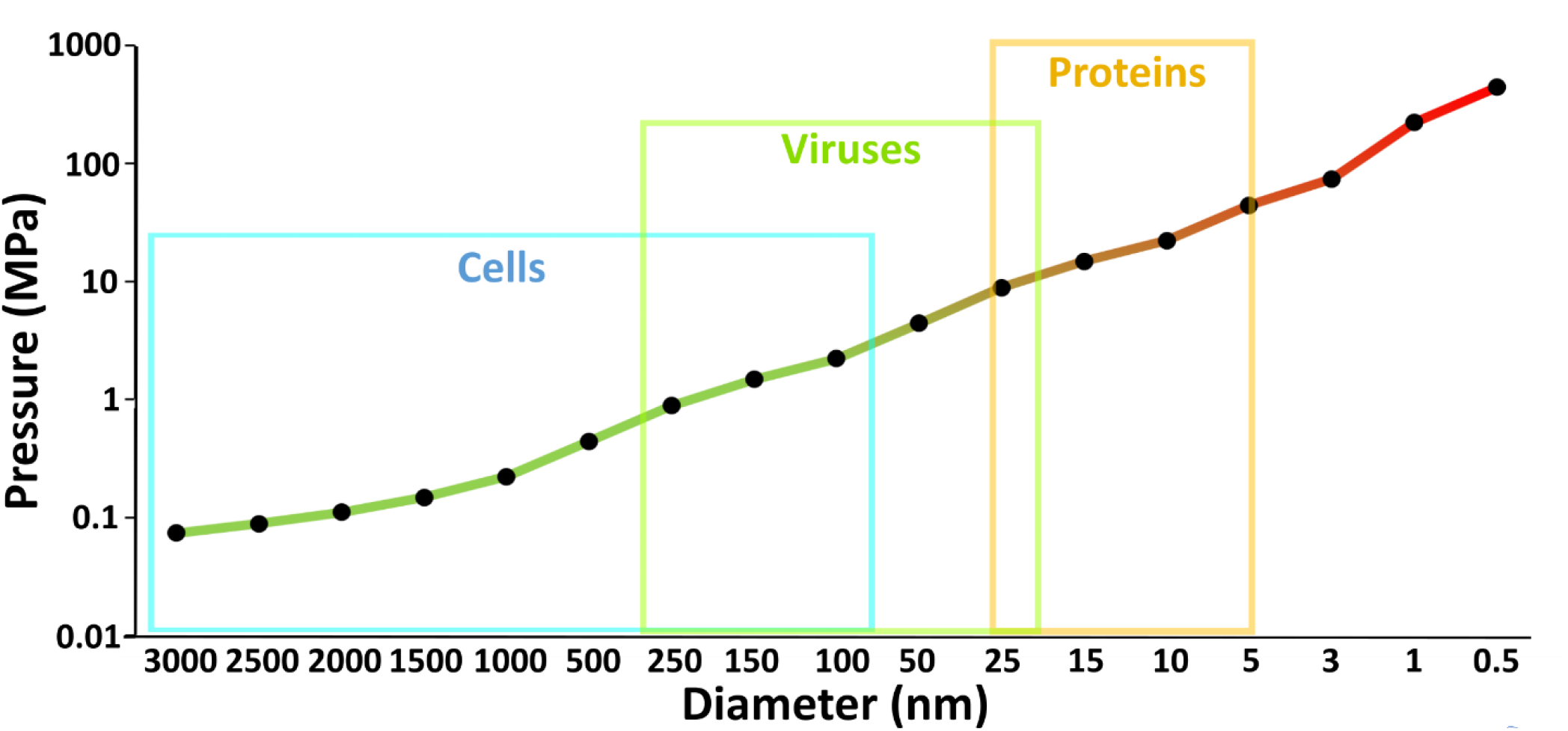
Analysis of the pressure on different volumes of spherical, graphene-encapsulated structures and its correlation with the size of relevant biological structures.

**Fig. S4.**
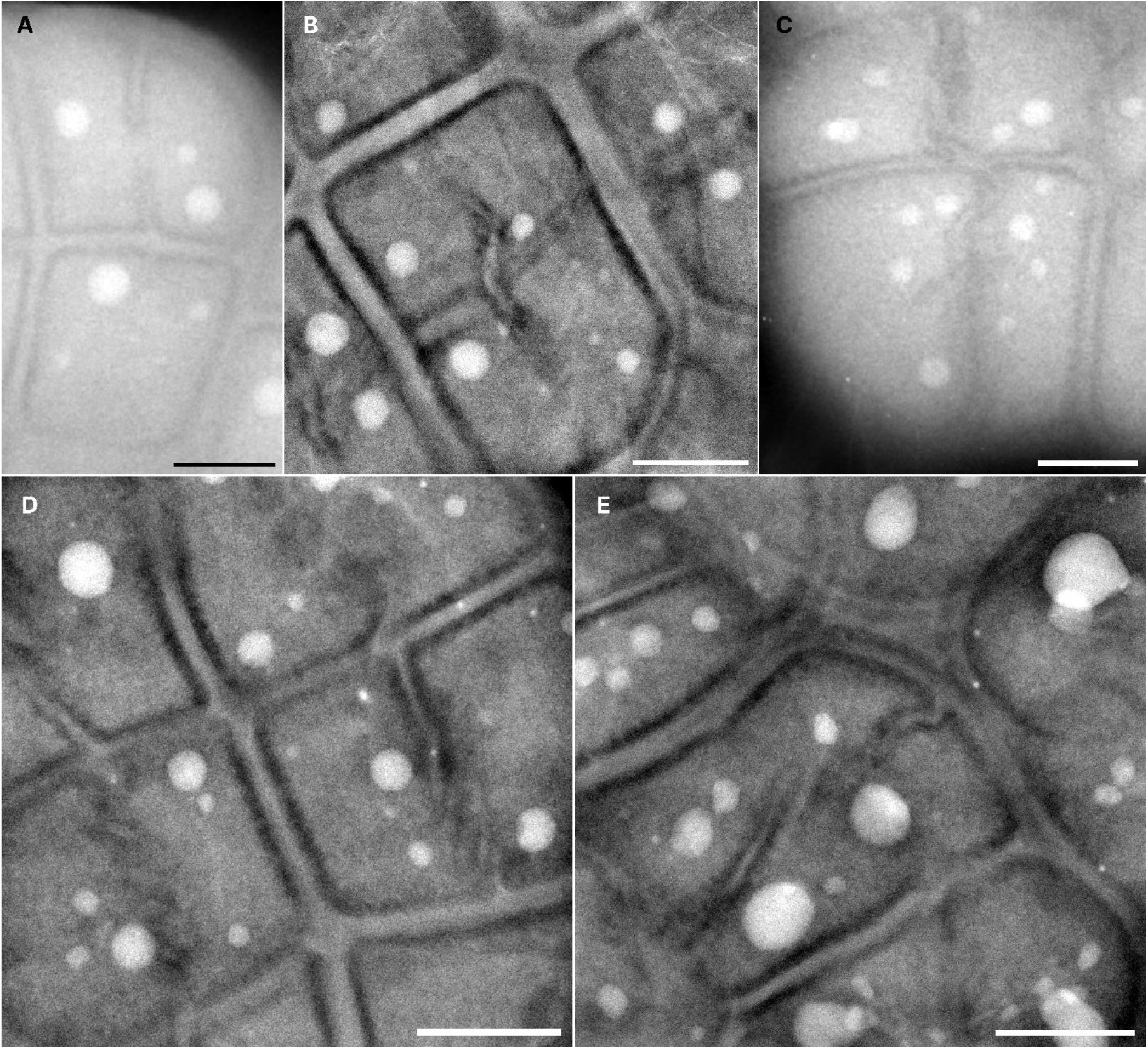
Asymmetric division and other irregularities in septal development in *D*.*radiodurans*. **A)** Individual cocci within a tetrad at different stages of cell division (Late stationary phase). **B)** S-shaped septum without complete perpendicular septum (Late stationary phase). **C)** S-shaped septum with complete perpendicular septum (Early exponential phase**). D)** Diagonal septa within individual cocci in late stationary phase and early exponential phase **(E)**. Scale bar: 500 nm.

**Fig. S5.**
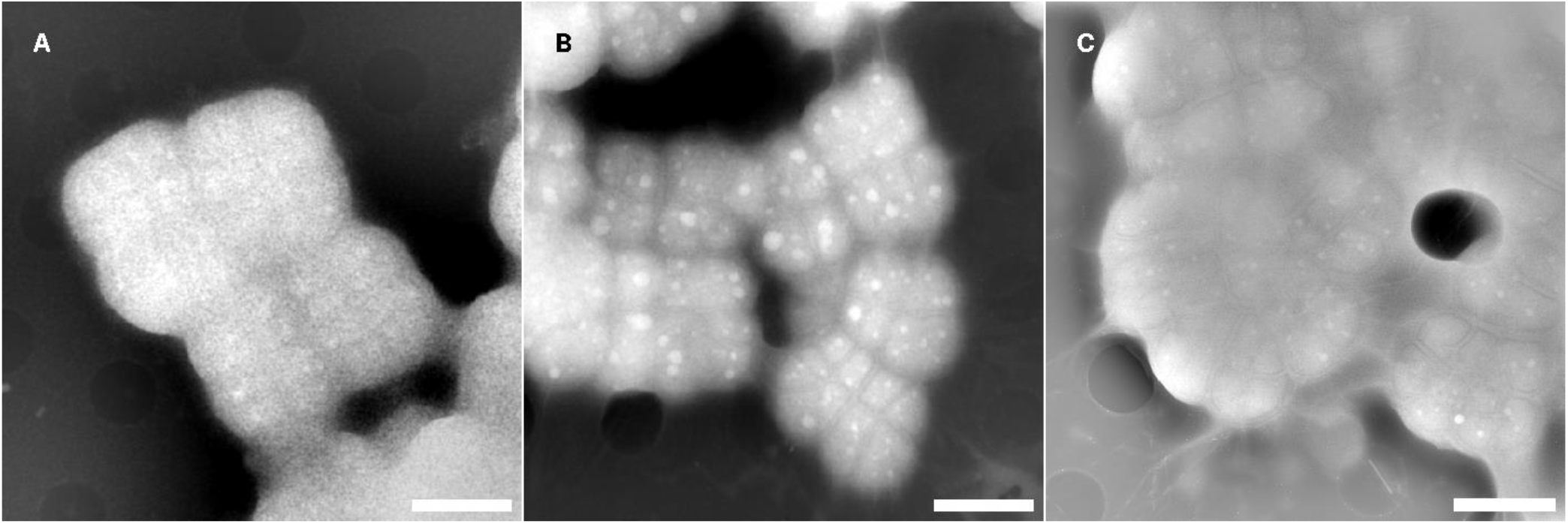
Imaging *D*.*radiodurans* at different fluence. BP-filtered LP-ADF-STEM images of *D*.*radiodurans* at fluences of **A)** 0.5, **B)** 5.8 and **C)** 65 e/nm^2^. Each individual coccus and endogenous polyphosphates are clearly visible at fluences as low as 5.8 e^-^/nm^2^. Scale bar: 2 μm.

**Fig. S6.**
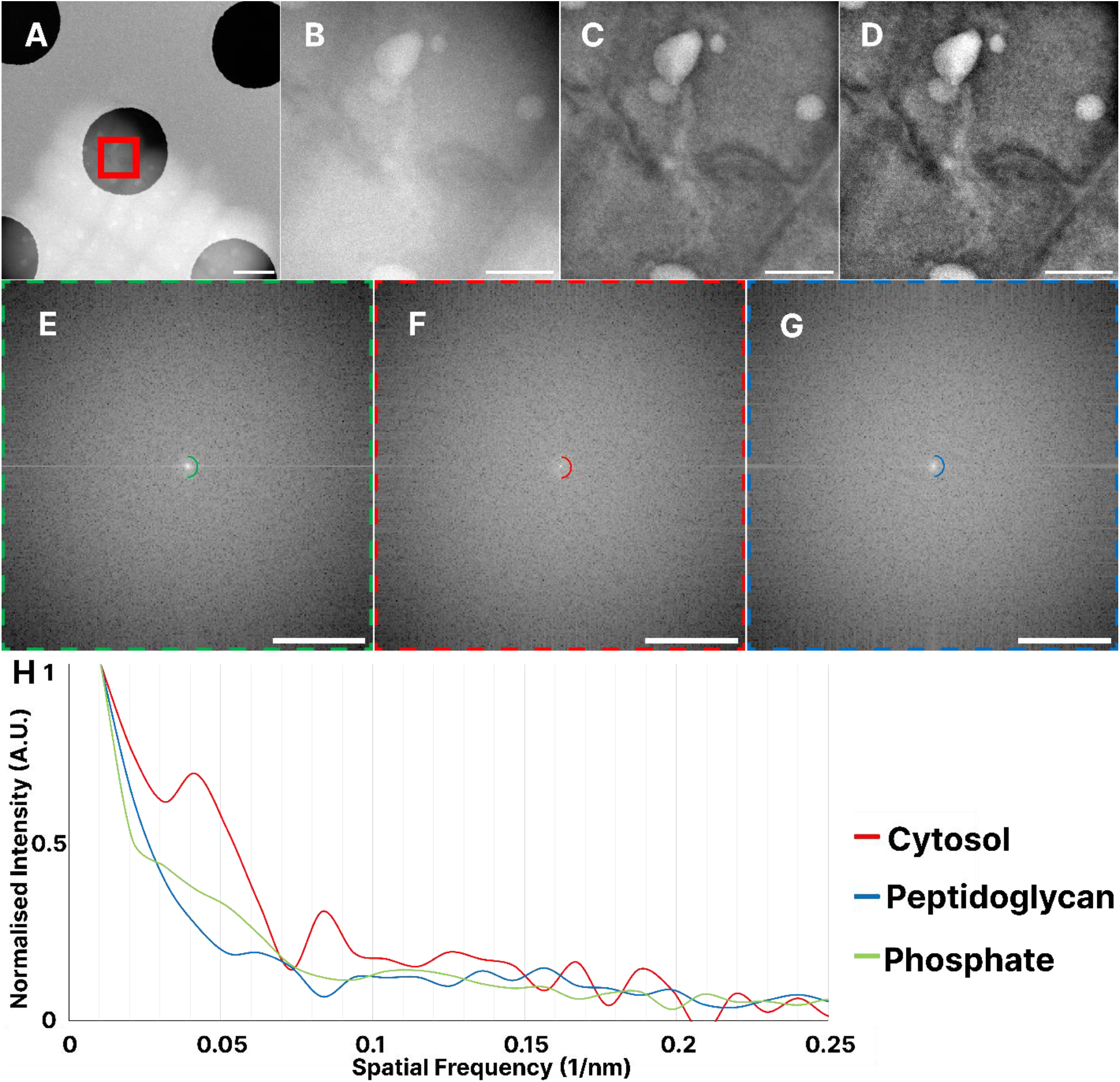
*D*.*radiodurans* ultrastructure in graphene encapsulated environments. **A)** Low magnification ADF-STEM image of encapsulated grouped *D*.*radiodurans* tetrad, Fig.3A ROI marked in red. **B)** Raw ADF-STEM image of *D*.*radiodurans*, **C)** After bandpass filter applied to **(B)**, after contrast enhancement to **(C)**. Corresponding full size power spectra for ROIs in Fig. 3B of storage granules **(E)**, cytosol **(F; Fig. 3B inset)** and peptidoglycan **(G)**, (18 nm)^-1^ boundary marked by semicircles. **H)** Radial profile plot of the power spectra **(E-G)**, demonstrating a peak frequency ∼0.0417 nm^-1^ in the cytosol not present in the peptidoglycan. Scale bar: A: 1 μm; B-D: 250 nm; PS: (2 nm)^-1^.

**Fig. S7.**
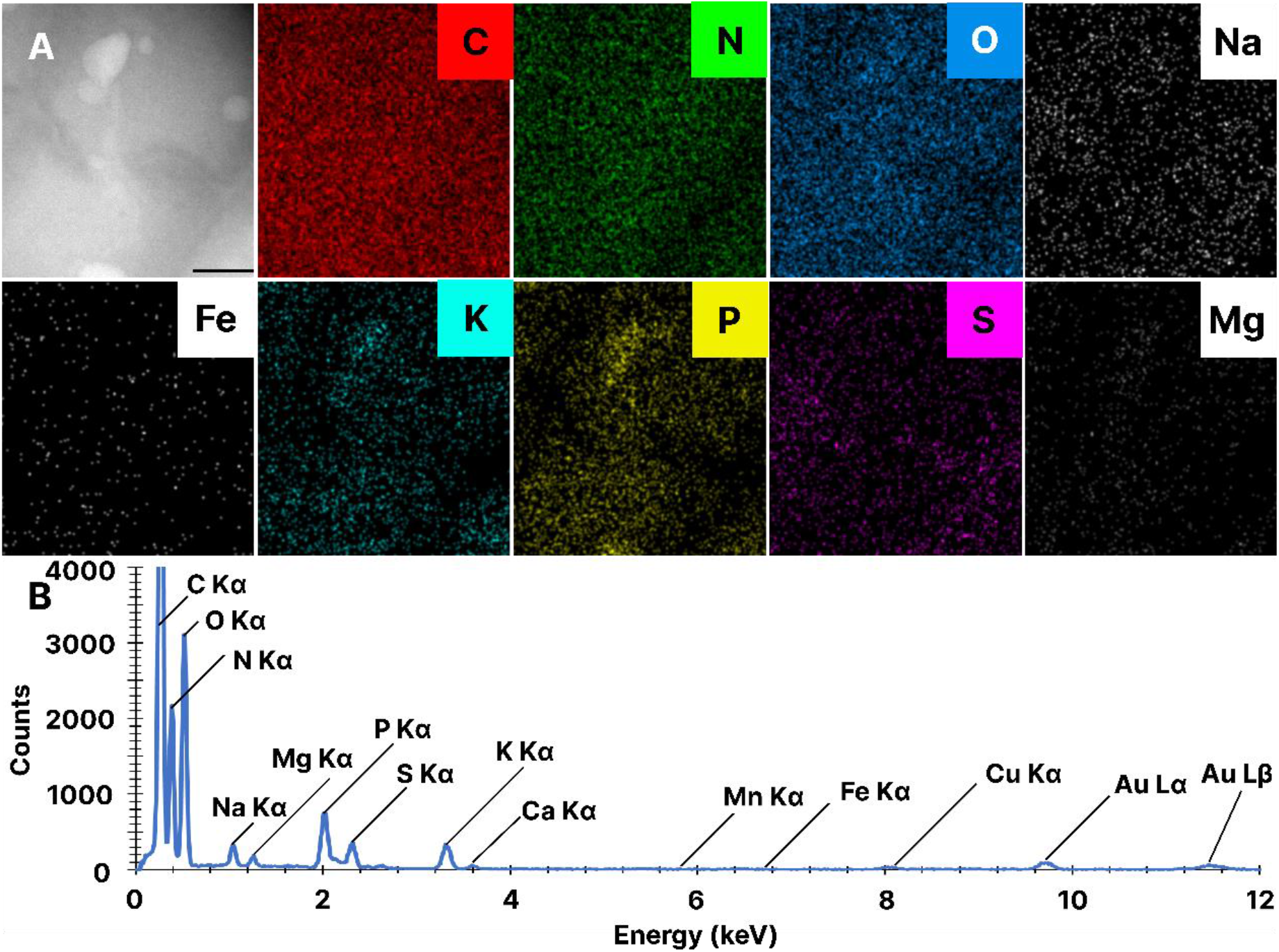
*D*.*radiodurans* elemental distribution in graphene encapsulated environments. **A)** Raw ADF-STEM image of Fig.3A and individual element maps. **I)** EDX spectrum of **(A)**, showing different elemental peaks, with no signal evident for Fe Kα lines. Scale bar: A: 250 nm.

**Fig. S8.**
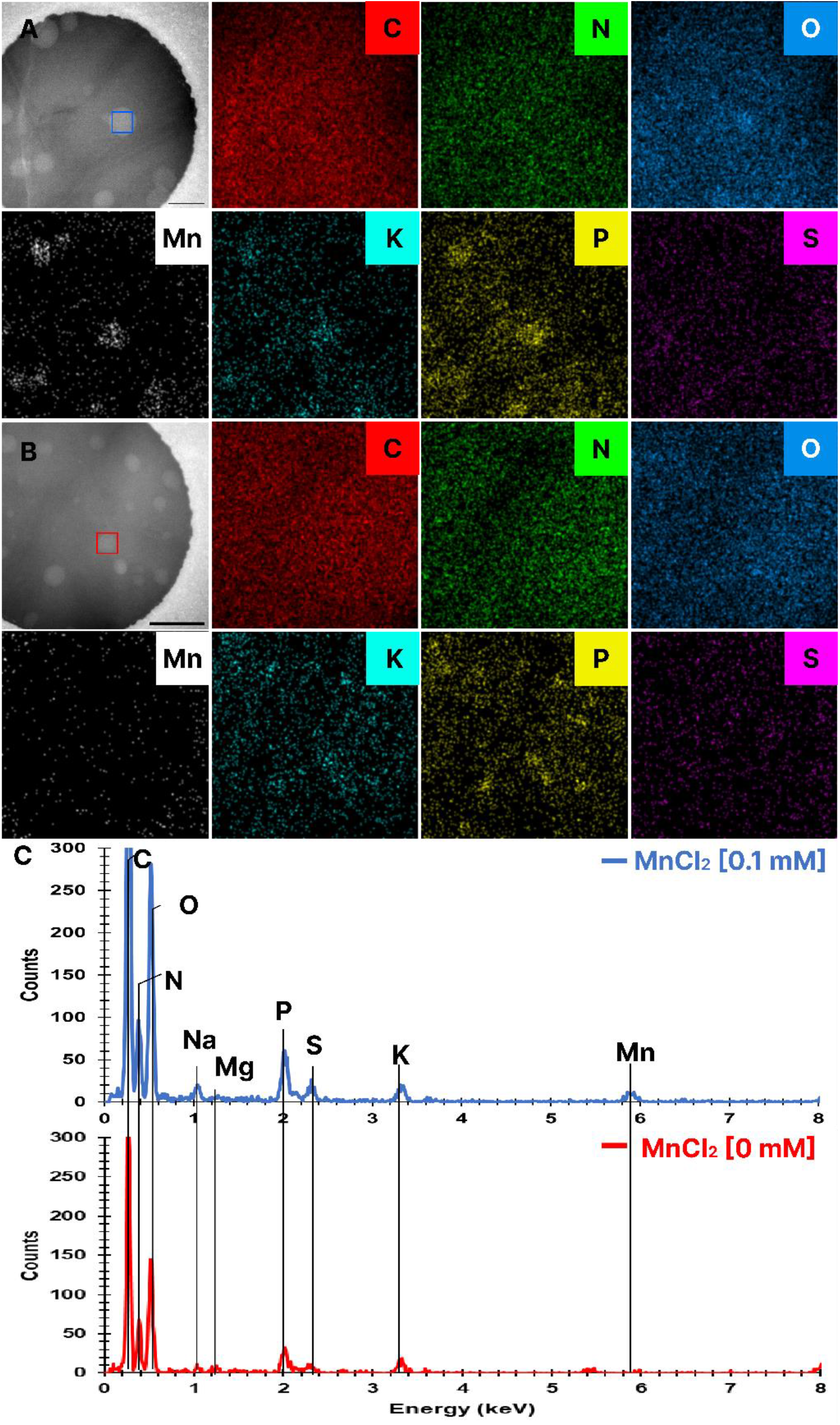
Manganese accumulation in bacterial phosphate granules. Individual element maps of D.radiodurans treated with **(A)** and without (**B)** MnCl_2_. **C)** EDX spectra of individual phosphate granules, selected area marked with dotted lines in **(A)** and **(B)**. Scale bar: A-B: 500 nm.

